# OocystMeter, a machine-learning algorithm to count and measure *Plasmodium* oocysts, reveals clustering patterns in the *Anopheles* midgut

**DOI:** 10.1101/2025.06.28.662088

**Authors:** Duo Peng, Eryney Marrogi, Elisabeth Nelson, Qiushi Liu, Tasneem A. Rinvee, Laura E. de Vries, Kate Thornburg, Naresh Singh, W. Robert Shaw, Flaminia Catteruccia

## Abstract

We present OocystMeter, a machine learning-based software developed to automate the segmentation of malaria oocysts from images of mosquito midguts stained with mercurochrome. Existing bioimage analysis tools, including machine learning-based ones, often struggle with the unique staining patterns, complex midgut backgrounds, and variable morphology of oocysts, making the determination of oocyst size and numbers cumbersome. To overcome these challenges, we curated a high-quality dataset comprised of 11,178 *Plasmodium falciparum* oocysts in *Anopheles gambiae* midguts annotated by expert parasitologists. Using this dataset, we fine-tuned a Mask R-CNN object detection model to achieve segmentation accuracy comparable to human parasitologists (Spearman’s correlation of 0.998 for oocyst counts and 0.978 for size measurements). Applying this tool in conjunction with spatial analysis, we uncovered a non-random, clustered spatial distribution of oocysts independent of the midgut’s anatomical regions or geometric axes, particularly in infections with fewer than 75 oocysts/midgut. Our workflow significantly accelerates malaria oocyst intensity and size analysis, reduces human bias, and provides spatial coordinates for advanced parasitology studies. OocystMeter is freely available at https://github.com/duopeng/OocystMeter, and as a web tool at http://Oocystmeter.org/, offering a valuable resource for researchers investigating the oocyst stage of malaria development.

## INTRODUCTION

Malaria transmission is strongly influenced by the number of oocysts in the midgut of the *Anopheles* mosquito vector, as well as by the speed of oocyst development [1–3]. After mosquitoes ingest *Plasmodium* gametocytes during a blood meal, they develop into gametes that fertilize to form a zygote. Shortly after, the zygote transforms into a motile ookinete that crosses the midgut wall and forms an oocyst underneath the basal lamina. Over a few days, the oocyst undergoes a spectacular process of growth and DNA replication which culminates with the formation of thousands of sporozoites, the form infectious to humans, that migrate to the mosquito’s salivary glands, ready for transmission.

Quantifying oocyst intensity and growth in the mosquito midgut is an important step in researching parasite biology and is also needed to evaluate the efficacy of novel malaria control interventions such as the use of anti-parasitic drugs in mosquito stages [4,5]. Antibody-based methods require time-consuming immunofluorescence staining [6], while protein-tagging involves genetically modified parasites, which often exhibit reduced transmission efficiency due to the fitness cost of transgenesis, and require fluorescence microscopes [7]. Near-infrared spectroscopy can detect parasite prevalence and intensity, but cannot measure oocyst size [8]. As a result, laboratories rely on staining oocyst-infected mosquito midguts with mercurochrome and using bright-field microscopy to assess oocyst number and size.

In this study, we developed an automated image-processing workflow for counting and measuring oocysts in mercurochrome-stained mosquito midguts. The workflow processes z-stack midgut images and returns oocyst counts, areas, coordinates, and masks using a mask R-CNN model [9] trained with samples from our laboratory. Using the spatial coordinates of oocysts in their midgut, we reveal a non-random and clustered pattern of oocyst distribution that is not driven by previously reported anterior-posterior enrichment [10,11] (ref. 10, not statistically significant). Our workflow accelerates oocyst intensity and growth kinetics analysis, significantly reducing processing time compared to manual counting and sizing, minimizing observer bias, and enabling precise acquisition of spatial coordinates for further investigation.

## RESULTS

### A machine learning algorithm trained to count and size oocysts

To train a machine learning model to count and size oocysts in mercurochrome-stained mosquito midguts, we created a training dataset comprising 11,178 oocysts collected from 231 images from midguts dissected at day 7–9 post infection. The distributions of per-midgut oocyst count and mean area are summarized in **Figure S1**. Oocysts were manually labeled with the VGG Image Annotator [12] (VIA) v2.0.7 using ellipses tracing their outline as drawn by trained parasitologists. We also labeled the outline of each midgut using the “polygon region shape” function.

From these training dataset, we first fine-tuned two bioimage analysis machine-learning models: StarDist [13] (designed for star-convex shapes, which closely resemble the morphology of oocysts) and the Cellpose [14] cyto2. To evaluate model performance, we conducted a threshold sweep of intersection over union (IoU) values between predicted segmentation masks and ground truth annotations to establish precise criteria for oocyst identification (**Figure 1a**). At an IoU threshold of 0.7, the StarDist model demonstrated an average precision (AP) of 71% (correctly identified oocysts relative to total predictions) and average recall (AR) of 60% (proportion of actual oocysts successfully detected). The Cellpose cyto2 model exhibited comparable performance, with AP of 68% and AR of 64% (**Figure 1a**). However, these performance metrics were inadequate for robust automated analysis, necessitating the evaluation of alternative models.

**Figure 1.**
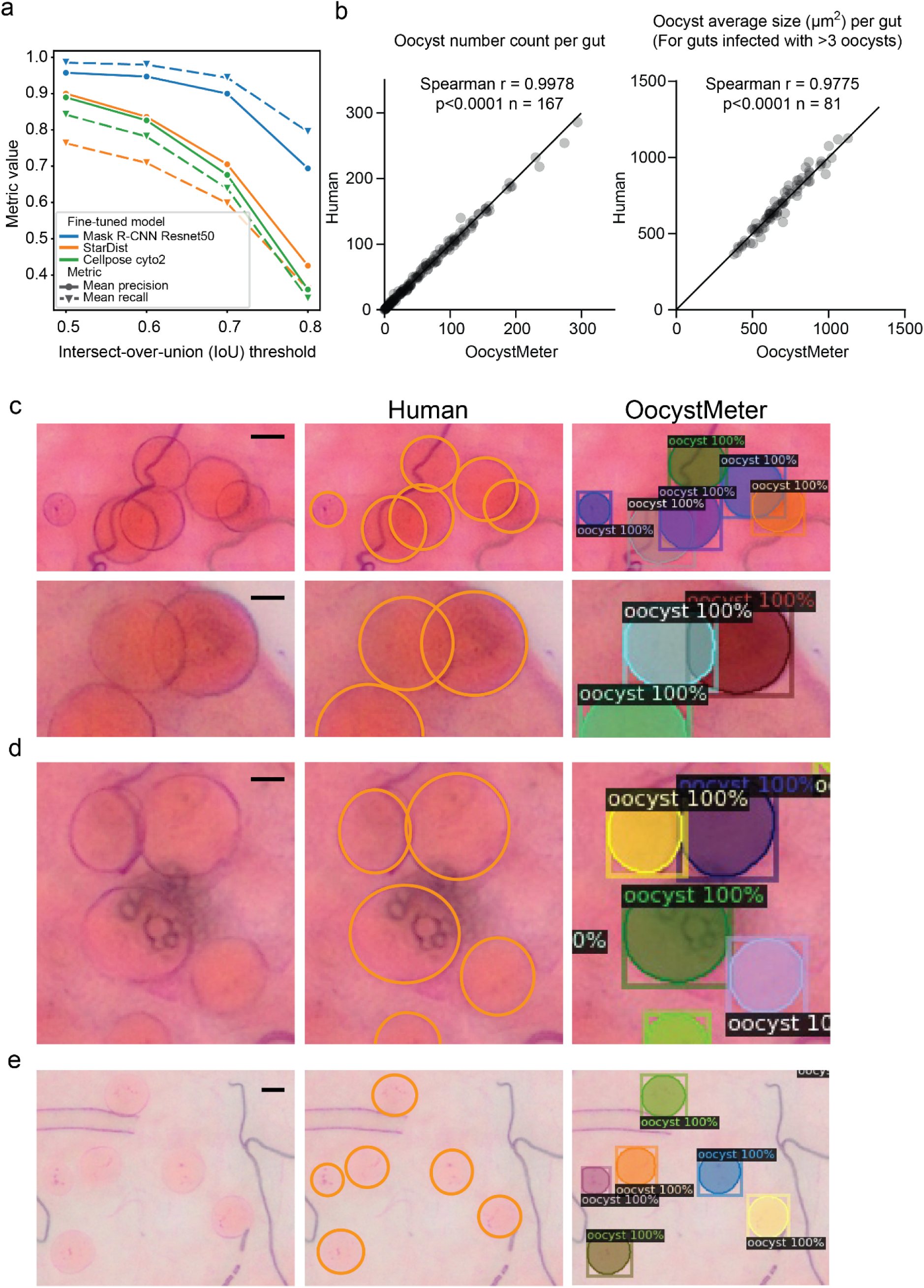
Performance of the oocyst segmentation OocystMeter model. **(a)** Average precision (AP) and average recall (AR) values under different thresholds of intersection over union (IoU) for different models. The Finetuned Mask R-CNN Resnet50 model is denoted as OocystMeter. **(b)** Oocyst number counts and average oocyst size matched closely between humans and the OocystMeter model. Each data point represents oocyst count or average oocyst size in a midgut. The diagonal line is drawn in solid black. Midguts with less than three oocysts were excluded from the analysis. See Figure S4 for the same analysis using midguts with fewer than four oocysts. **(c-e)** The OocystMeter model accurately performs segmentation with **(c)** overlapping, **(d)** partially obscured, and **(e)** weakly stained oocysts. Detection bounding boxes, confidence, and segmentation masks are shown for OocystMeter model results. Scale bars represent 10 μm. In all panels, midguts were dissected at 7–9 days post infection.

Next, we fine-tuned a more general object detector model: Mask R-CNN, which is pre-trained on the publicly available Common Objects In Context (COCO) dataset [15]. We hypothesized that a general object detection model would provide broader image processing and feature extraction capabilities, serving as a stronger foundation for fine-tuning into a highly specialized model. The fine-tuned Mask R-CNN model, hereafter referred to as the OocystMeter model, achieved 94.4% AP and 90.0% AR using a threshold IoU of 0.7 (**Figure 1a, Table S1a**). The OocystMeter’s performance exceeded that of fine-tuned StarDist and Cellpose cyto2 models, while achieving results comparable to those obtained by parasitologists. Oocyst counts and average oocyst size per midgut matched closely between the OocystMeter model and humans, reaching a Spearman’s correlation coefficient of 0.9978 and 0.9775, respectively (**Figure 1b**). This small discrepancy in average oocyst size (**Figure 1b, right**) is comparable to the discrepancy observed between two humans (**Figure S2**). The OocystMeter model performed well with overlapping (**Figure 1c**), partially obscured (**Figure 1d**), and weakly stained (**Figure 1e**) oocysts. The trained weights of the model and computer code required to perform oocyst counting and size measurement are available at: https://github.com/duopeng/OocystMeter.

### Full automation of the image analysis workflow

To further automate the oocyst analysis, we developed an image preprocessing pipeline as part of the workflow to transform multiple z-stacks into a single 2D view of a midgut, imaged at 100X or 200X magnification (**Figure 2**). The pipeline first uses ImageJ macros to flatten z-stacks of two or more fields of view, then stitches together flattened z-projections based on overlapping regions, creating a single panoramic view covering the entire midgut for the OocystMeter model to analyze. To put oocysts in spatial context and facilitate downstream spatial statistics analysis, we created a midgut model to segment the boundary of midguts (**Figure 2**, lower left). This pipeline can preprocess a typical midgut (two regions with z-stacks of two images each) in under 1 minute on a personal computer without the need for GPUs or specialized computing hardware. The fully automated pipeline is available publicly as a web tool service at http://Oocystmeter.org. The computer code for the image preprocessing pipeline is available at https://github.com/duopeng/image_merge-z-stack_and_stitch.

**Figure 2.**
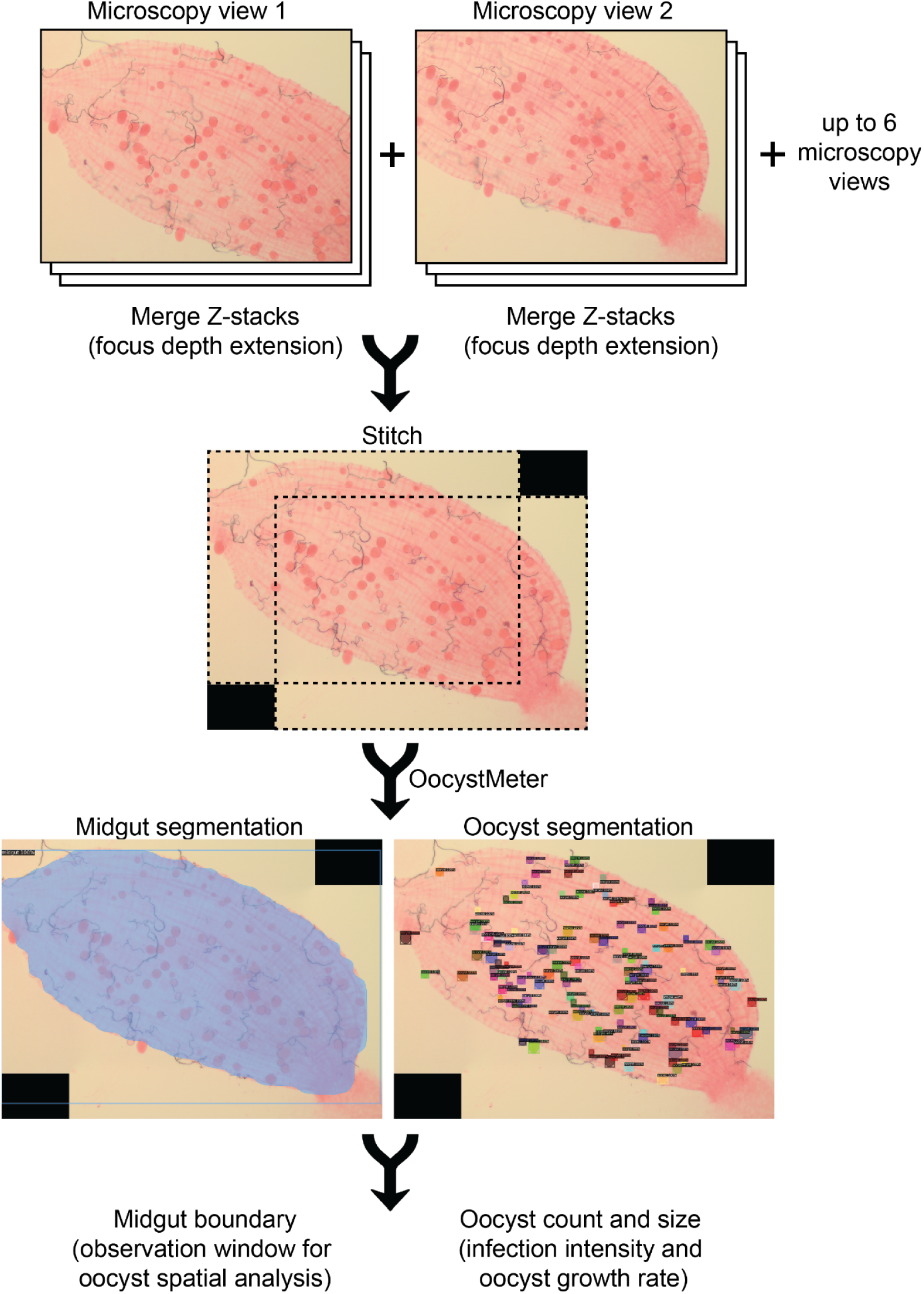
Automated oocyst analysis workflow. The workflow first performs focus depth extension by merging two (or more) z-stacks of images taken at the same field of view, then up to six fields of view can be stitched to form a panorama view covering the whole midgut. The panorama view is then given to the OocystMeter model to count oocysts and measure their sizes. The output also includes the boundary of the midgut, which could be used as the observation window in oocyst spatial distribution analysis.

### Oocysts are non-randomly distributed in the midgut

An additional output of the OocystMeter tool is the digital coordinates of oocysts relative to the midgut perimeter. Using these data, we analyzed the spatial distribution of oocysts within the midgut to determine whether it deviates from complete spatial randomness (CSR), also known as the spatial Poisson process [16]. To this end, we represented each oocyst as a dimensionless point at its centroid in a Cartesian coordinate system. The midgut boundaries were converted into polygons within the same coordinate system to serve as observation windows.

We stratified our midgut images into oocyst intensity categories: low (4 ≤ n ≤ 25), medium (26 ≤ n ≤ 74), and high (n ≥ 75) (**Figure 3a**). Notably, these thresholds—especially for the high-intensity category—are considerably higher than those typically observed in natural infections [17], where infections with over 25 oocysts are generally classified as high intensity.

**Figure 3.**
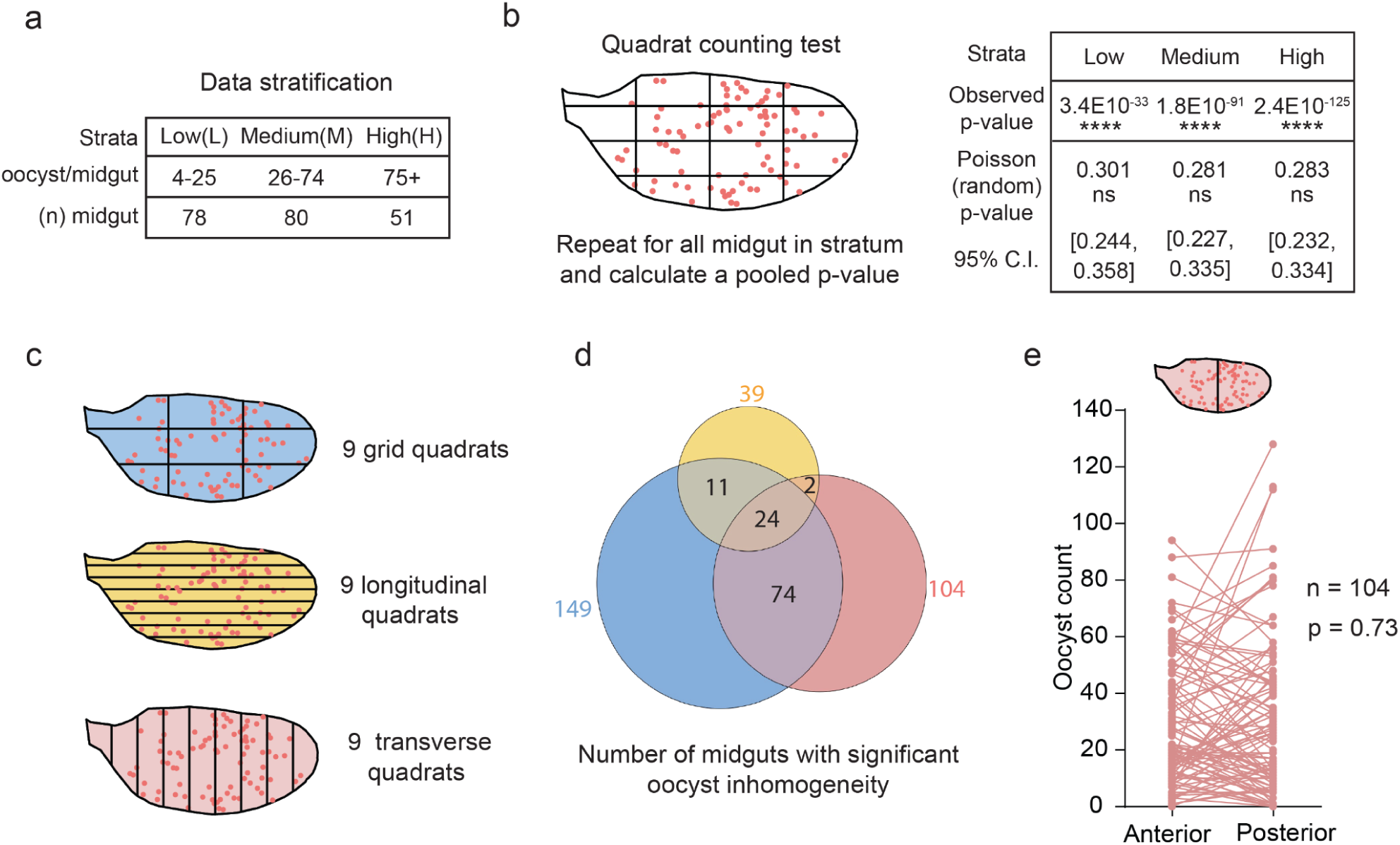
The distribution of oocysts in the midgut is not random. **(a)** Midgut images are divided into low, medium, and high infection intensity categories. **(b)** The quadrat counting test indicated that all categories had a statistically significant inhomogeneous distribution of oocysts. Each midgut was divided into 16 quadrants (grid-style), and the quadrant count statistics of all midguts in each category were pooled to calculate a single p-value. For each category, 1,000 Poisson simulations were repeatedly performed on each midgut with a matching number of oocysts, and the 95% confidence intervals were calculated. **(c-e)** Oocyst inhomogeneity is not tied to a single anatomical axis of the midgut. **(c)** Grid, longitudinal, and transverse quadrat partitioning schemes. **(d)** Venn diagram showing the midguts displaying inhomogeneous oocyst distribution in three different quadrat partitioning schemes. Numbers next to each primary circle indicate the number of midguts significant for the corresponding quadrat partitioning schemes. Numbers in overlapping regions indicate midguts that are significant for multiple quadrat partitioning schemes (significance level = 0.05, each midgut was analyzed independently by each quadrat type). **(e)** There is no enrichment of oocysts in the anterior or posterior halves of the midgut. Midguts exhibiting inhomogeneity along the anterior-posterior (n = 104) were analyzed (Wilcoxon matched-pairs signed-rank test, p = 0.73).

Images within the same infection intensity category were treated as repeated observations of the same spatial distribution process. We first conducted a quadrat counting test of CSR (**Figure 3b, left**), which resulted in significant p-values for all three categories, indicating a strong rejection of CSR (**Figure 3b, right**). This finding was confirmed by randomly placing the same number of oocysts in each midgut according to a homogeneous Poisson point process (equivalent to CSR) and repeating the simulation 1,000 times. The mean p-value and the 95% confidence interval from the simulations did not overlap with the observed p-value (**Figure 3b, right**), reinforcing the rejection of CSR and supporting the alternative hypothesis that oocysts are not randomly distributed in the midgut.

### Oocyst inhomogeneity is not tied to a single anatomical axis of the midgut

To further investigate the factors contributing to the non-random distribution of oocysts, we examined whether spatial gradients along the anterior-posterior or dorsal-ventral axes of the midgut could explain oocyst enrichment. Previous studies have reported oocyst enrichment in the posterior of the midgut for two parasite-mosquito combinations, and this trend was not influenced by mosquito orientation [10,11] (ref. 10, not statistically significant). For instance, in *Aedes aegypti* infected with *Plasmodium gallinaceum*, the posterior half of the midgut had significantly more parasites than the anterior half in both control mosquitoes and mosquitoes kept in continuously rotating containers [11].

To this end, we extended the classical quadrat counting test by implementing longitudinal and transversal quadrat schemes (yellow and salmon colors, respectively, **Figure 3c**). In these schemes, longitudinal quadrats partition the midgut with lines parallel to the anterior-posterior axis, while transversal quadrats use lines perpendicular to this axis. Each midgut was independently analyzed using all three quadrat-partitioning schemes. Of the 149 midguts with significant oocyst inhomogeneity by the classical quadrat-counting test (blue, **Figure 3c,d**), 98 were also significant in the transverse quadrat analysis (overlap of the blue and salmon circles in the Venn diagram, **Figure 3d**), 35 in the longitudinal analysis (overlap of the blue and yellow circles, **Figure 3d**), and 24 in both. These patterns indicate that oocyst inhomogeneity is not tied to a single anatomical axis of the midgut.

For the 104 midguts that exhibited inhomogeneity along the anterior–posterior axis (salmon color, **Figure 3c,d**), we compared oocyst counts in the anterior and posterior halves using a Wilcoxon matched-pairs signed-rank test and found no significant difference (**Figure 3e**).

### Oocyst distribution follows a clustering pattern

For a more granular analysis of the oocyst distribution pattern, we performed Ripley’s K function analysis. This analysis is more granular in two ways: (a) it analyzes spatial patterns as a function of analysis radius (r); and (b) it can distinguish two types of non-randomness: clustering (can be caused by attraction, patches of favorable location and synergistic interaction), and regular spacing (can be caused by avoidance or local inhibition). To construct randomness baselines, we employed two simulation approaches: (i) a homogeneous Poisson point process which generates randomly distributed points in each corresponding midgut observation window, with number of points matching the number of oocysts (**Figure 4a top left, random baseline 1**); and (ii) random points sampled from the surface of an ellipsoid and then projected onto a 2D plane, simulating the flattening of a midgut from its 3D ellipsoid-like structure (**Figure 4a right, random baseline 2**).

**Figure 4.**
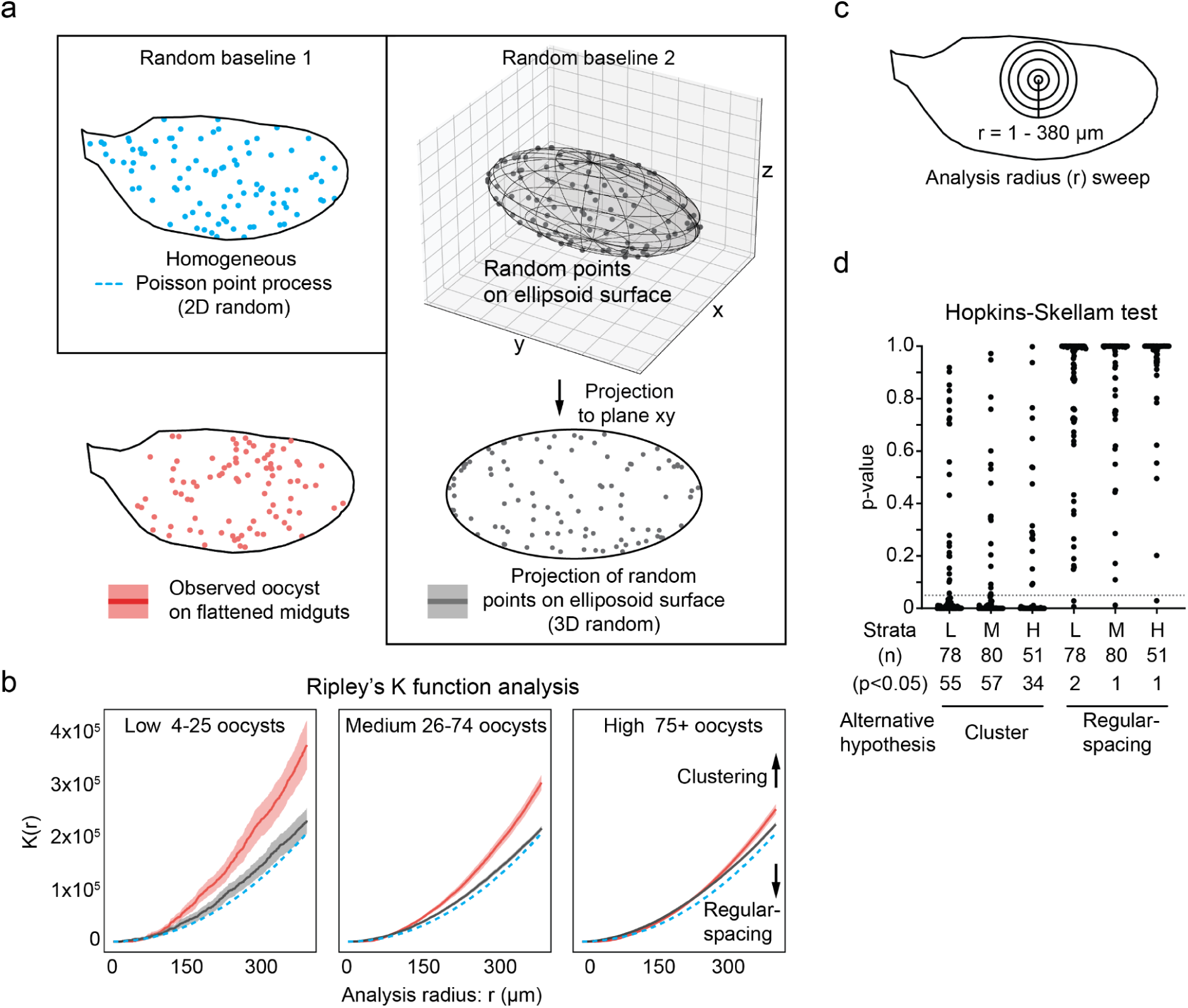
Oocyst distribution follows a clustering pattern. **(a-c)** Ripley’s K function analysis indicated that all infection intensity categories had a significant clustering pattern of oocyst distribution. (**a**) The homogeneous Poisson point process was used to generate random oocysts in the matching midgut area (top left, Random baseline 1). To simulate the process of random oocysts on the 3D midgut surface being compressed to a 2D space during microscopy, random points were sampled from the surface of an ellipsoid and projected onto a 2D plane (Top right, random baseline 2). When generating random baselines, the number of repeated observations matched midgut numbers in each infection intensity category, and the number of data points in each observation matched the oocyst number in the corresponding midgut. Each midgut was treated as a replication of the same biological process in the category. A 95% confidence interval was shown for the real oocyst data and 3D surface random reference, and the 95% confidence interval for Poisson simulations is extremely narrow and indistinguishable from the dashed line. **(b)** Ripley’s K function, K(r), was calculated for three different infection intensity strata and their corresponding random baseline controls. Note that the wider 95% confidence interval in the low infection stratum is due to the low sampling power caused by a low number of data points (oocysts) in each midgut. **(c)** Schematic drawing showing K(r)’s analysis radius (r) with respect to a typical midgut size. **(d)** The Hopkins Skellam test was performed with “Cluster” or “Regular-spacing” alternative hypotheses (HA), with the null hypothesis set to a homogeneous Poisson point process. Each midgut generates a single p-value for each HA (significance level = 0.05).

We calculated Ripley’s K function for each midgut image and generated an average K(r) curve and 95% confidence interval for each stratum (**Figure 4b**, red line with ribbon). Ripley’s K function calculated for radii (r) from 0–380 µm (**Figure 4c**) deviates significantly from both a homogeneous Poisson and a 3D random process (**Figure 4b**, blue dotted lines and solid gray lines with ribbon, respectively). Our data reveal a statistically significant departure from randomness across all three categories of infection intensity. The K(r) line and its 95% confidence interval band (red) remained above the two random references (blue dotted lines and solid gray lines with ribbon), suggesting a statistically significant clustering pattern as opposed to a regular spacing pattern (**Figure 4b)**. For the low and medium oocyst number category (4–25 and 26–74 oocysts per midgut, respectively), the clustering pattern is observable with an analysis radius (r) of >150 µm (about three oocysts across), while for the high oocyst number category (75 and more) the clustering pattern is only observed when the analysis radius (r) >300 µm (∼ 6 oocysts across).

To confirm the clustering pattern, we conducted a Hopkins-Skellam test in which alternative hypotheses can be defined as clustering or regular spacing patterns. A multi-test-corrected p-value was generated for each image, testing for two alternative hypotheses (**Figure 4d**). In the low and medium oocyst stratum, 55 out of 78 and 57 out of 80 midgut images show significant evidence of a clustering pattern, respectively (**Figure 4d**). In the high-oocyst stratum, 34 out of 51 midgut images show significant evidence of a clustering pattern (**Figure 4d**). On the other hand, very few (n=4) midgut images tested significantly for a regular pattern of oocyst distributions, thus confirming that oocyst distribution mostly follows a clustering pattern. The Clark-Evans test, which detects deviations from CSR by analyzing neighbors of each point (as opposed to analyzing random pairs of points in the Hopkins-Skellam test), showed a similar result (**Figure S3**). Together, these additional statistical tests confirm our observation that the majority of infected guts show an oocyst clustering pattern.

## DISCUSSION

In this study, we present a machine learning software designed to identify and measure the size of *P. falciparum* oocysts in *An. gambiae* midguts stained with mercurochrome. Our tool generates oocyst counts and size measurements that align with those produced by trained parasitologists, but in roughly one-tenth the time (**Figure 1b**). The time savings is more pronounced in heavy infections. Our machine learning model was trained on oocyst annotations aggregated from four parasitologists, averaging out individual biases that often arise when handling ambiguous cases, such as obscured, burst, or faintly stained oocysts. Our tool ensures objective and consistent decisions in these instances. The inclusion of highly infected midguts in the training data enabled the software to accurately differentiate overlapping oocysts and comprehensively identify hundreds of oocysts in a single midgut (up to 500 oocysts per midgut). A caveat is that this tool was optimized for oocysts at 7–9 days post infection, when oocysts are 500–1500 μm^2^ in size and more easily identified. Detection sensitivity decreases for very small oocysts (<200 μm^2^) in earlier developmental stages, which may result in slight undercounting during the initial days post-infection. We believe this tool is useful for quantifying *P. falciparum* oocysts across diverse research settings, and can be readily adapted to other *Plasmodium*-*Anopheles* combinations, such as *P. berghei* in *An. stephensi*, after validation and/or further fine-tuning with species-specific images.

The oocyst growth rate, typically estimated by measuring oocyst size at multiple time points following an infectious blood meal, is a crucial metric in studying the complex interactions between the oocyst and its mosquito host, including but not limited to immunological responses and nutrient exchange (as reviewed by Shaw et al. [18]). By removing significant labor barriers, our tool paves the way for accelerated research into factors influencing oocyst growth, screening for oocyst-targeting drugs, and the development of novel strategies to curb malaria transmission [5,19].

Capitalizing on our rich dataset of oocyst spatial coordinates relative to the midgut boundary, we have documented a non-random distribution of oocysts in the mosquito midgut. The non-random distribution is predominantly a clustering pattern, as opposed to a regular spacing pattern. In low and medium infections (4–25 and 26–74 oocysts per midgut, respectively), the oocyst clustering pattern was detectable from an analysis radius of 150 µm and above, while for the high infection category, the oocyst clustering was detectable with an analysis radius of above 300 µm (**Figure 4b**). A possible explanation for these data is that high infection intensities result in overcrowding of oocysts, obscuring local clustering patterns. Additionally, parasite-mosquito interactions may be significantly different in high-intensity infections. For example, large numbers of traversing ookinetes may cause excessive tissue damage and cell apoptosis of the midgut epithelium, disrupting the underlying mechanisms that generate the “clustering force”.

Several possible factors could exert a “clustering force” on oocyst: (i) microenvironment created/maintained by the invasion process itself or regional midgut cells, (ii) midgut regional features (e.g., trachea, innervation), and (iii) gravity or other physical factors. Prior research showed evidence of enrichment of *P. berghei* and *P. gallinaceum* oocysts in the posterior midgut of *An. stephensi* [10] and *Ae. aegypti* [11], respectively. However, in this study, we did not observe *P. falciparum* oocysts enriched in the posterior or anterior midgut of *An. gambiae* (**Figure 3e**). Further studies are needed to determine the biological factors influencing the non-random, clustered spatial distribution of oocysts along the basal lamina of the midgut. Clarifying these drivers may enrich our understanding of oocyst biology and direct the design of mosquito-targeted interventions to interrupt malaria transmission.

## MATERIAL AND METHODS

### Rearing of *Anopheles gambiae* mosquitoes

The *An. gambiae* wild-type G3 strain was reared in an insectary, and the climate was maintained at 26–28 °C, 65–80% relative humidity, 12:12 hr light/darkness photoperiod. Adults in colony cages were fed on 10% glucose solution *ad libitum*. Virgin female mosquitoes were obtained by separating the sexes at the pupal stage after a visual examination of the terminalia.

### Culturing of *Plasmodium falciparum* NF54 parasites

*P. falciparum* parasites (NF54 strain) were maintained in continuous culture using established methods [22,23]. Asexual parasite stages were grown in human red blood cells (Interstate Blood Bank, Memphis, TN) at 5% hematocrit, with parasitemia levels kept between 0.2–2%. Cultures were incubated at 37°C in RPMI 1640 medium containing 25mM HEPES, 10mg/l hypoxanthine, 0.2% sodium bicarbonate, and 10% heat-inactivated human serum (Interstate Blood Bank). A controlled gas atmosphere of 5% oxygen, 5% carbon dioxide, and balanced nitrogen was maintained throughout the culture period, which extended up to 8 weeks.

To produce gametocytes, parasitemia was increased above 4% and cultures were maintained for 14–20 days with daily medium changes. This extended incubation period allowed for the accumulation of mature stage IV and V male and female gametocytes.

### *P. falciparum* infection and imaging of infected midguts

Five-day-old female mosquitoes were transferred to a sealed, secure infection glovebox and provided with an *in vitro* culture of *P. falciparum* (NF54) gametocytes through a custom-made, glass, water-heated membrane feeder. After 60 min, female mosquitoes that failed to engorge fully were vacuum-aspirated out of their cages directly into 80% ethanol and discarded. At 7–9 days after an infectious blood meal, female mosquitoes were vacuum-aspirated into 80% ethanol, incubated for 10 min at −20 °C, and transferred out of the secure infection box into PBS on ice. Midguts were dissected out in PBS and stained with 0.2% w/v mercurochrome (in ddH_2_O) for 12 min. After staining, midguts were mounted on glass microscope slides in 0.02% w/v mercurochrome. Images of stained midguts were taken at 100X or 200X on an Olympus Inverted CKX41 microscope. The majority of the midguts were covered by two microscope views with two focal planes per view.

### Model training and validation

Midgut images were preprocessed with the focus-stacking (merge z-stack) and stitching pipeline developed in this study (available at https://github.com/duopeng/image_merge-z-stack_and_stitch). Oocysts in each midgut image were annotated with the ellipse shape drawing tool in the VGG Image Annotator (VIA) tool [12]. Midguts were annotated with the polygon drawing tool in VIA. Annotations were exported using the COCO format [15]. Images and corresponding annotations were split using an 80/20 ratio for training and validation purposes, respectively. Images were resized so that the long and short edges did not exceed 2300 and 1500 pixels, respectively. Training and validation datasets were registered with the Detectron2 [24] framework using author-provided functions. Hyperparameter tuning was performed based on empirical experience, and our best model (v3) was trained with the following non-default parameters:

model_network = COCO-InstanceSegmentation/mask_rcnn_R_50_FPN_3x.yaml;

loss = smooth_l1;

cfg.SOLVER.IMS_PER_BATCH = 2;

cfg.MODEL.ROI_HEADS.POSITIVE_FRACTION = 1;

cfg.DATASETS.PRECOMPUTED_PROPOSAL_TOPK_TEST = 4000;

cfg.DATASETS.PRECOMPUTED_PROPOSAL_TOPK_TRAIN = 4000;

cfg.MODEL.RETINANET.TOPK_CANDIDATES_TEST = 3500;

cfg.MODEL.RPN.POSITIVE_FRACTION = 1;

cfg.MODEL.RPN.POST_NMS_TOPK_TEST = 4000;

cfg.MODEL.RPN.POST_NMS_TOPK_TRAIN = 4000;

cfg.MODEL.RPN.PRE_NMS_TOPK_TEST = 4000;

cfg.MODEL.RPN.PRE_NMS_TOPK_TRAIN = 4000;

Our customized modifications of the detectron2 framework include: random image augmentation, a custom implementation of the evaluation of validation loss, and modification of the coco_eval.summarize2() function in pycocotools package (used by Detectron2) to increase the object instance detection limit to 500 (default=100). Code for our customized modifications can be found at https://github.com/duopeng/detectron2_customization.

The OocystMeter model was trained for 10000 iterations on a Tesla T4 or P100 GPU with 16 GB of GPU memory. Weights of the model were saved every 200 iterations, and the best weights were determined by examining the validation loss curve.

To enhance AP and AR for oocyst segmentation, we first expanded our training dataset. Utilizing the initial, less accurate OocystMeter model (v1), we accelerated this expansion. Model v1 pre-identified oocysts in new midgut images, which were then refined by two parasitologists to correct any inaccuracies in the oocyst masks generated by the model. This process added 145 images with 5725 oocysts to our dataset (**Table S1a**). Subsequently, we developed an improved OocystMeter model (v2) using this augmented dataset. Model v2 achieved an AP of 88.7%; however, the AR remained unchanged.

We initially observed that our OocystMeter model struggled to identify all oocysts in scenarios of high-intensity infections (exceeding 100 oocysts/midgut) or when oocysts were densely clustered. To address this, we adjusted two key hyperparameters in the region proposal network of the model: the number of proposed regions and the proportion of these regions considered ’positive’ for subsequent evaluation (positive fraction). Our hypothesis was that images with a dense oocyst population necessitate a larger number of proposed regions for initial detection, along with a higher proportion of these regions undergoing detailed evaluation. By increasing both the number of proposed regions and the proportion evaluated as potentially containing oocysts, our final OocystMeter model iteration (v3) achieved an improved AP of 94.4% and AR of 90.0% (**Figure 1a, Table S1a**).

### Spatial statistical analysis

The coordinates of oocysts and polygon boundaries of midguts were exported from VIA annotations and converted to point pattern datasets (ppp) implemented by the spatstat [16] package in the R programming language. Oocysts were treated as sizeless points, and the midgut boundaries were treated as observation windows. All the functions listed below were implemented by the spatstat package. Quadrat counting tests were performed with the quadrat.test function in spatstat. A 4x4 grid was specified by quadrat.test with arguments (ppp, 4, 4), and a 3x3 grid was specified by quadrat.test with arguments (ppp, 3, 3), nine transverse grids were specified by quadrat.test with arguments (ppp, 9, 1), nine longitudinal grids were specified by quadrat.test with arguments (ppp, 1, 9). Ripley’s K function analyses were performed by Kest with arguments (ppp, ratio=TRUE). K function analyses and quadrat count test results were pooled (stratum-wise) using the do.call(pool) function. Clark-Evans tests were performed by the clarkevans.test() function using the “guard” boundary correct option and “regular” or “clustered” as the alternative hypothesis. Clark-Evans indices were calculated by the clarkevans() function using the “guard” boundary correct option. The anterior and posterior oocyst counts were obtained by summing the anterior or posterior transverse quadrats, respectively.

## ACKNOWLEDGEMENTS

We thank Emily Selland for help with mosquito rearing, and members of the Catteruccia laboratory for comments on the manuscript. Funding for this study was provided by the NIH (R01AI148646; R01AI153404) to F.C.. D.P. was supported by the Harvard Data Science Initiative (Postdoctoral Fellow Research Fund, 2019), L.E.D.V. was supported by a Rubicon grant from the Dutch Research Council (NWO; grant number: 452021309). The findings and conclusions within this publication are those of the authors and do not necessarily reflect positions or policies of the HHMI, or the NIH. F.C. is an HHMI Investigator.

## AUTHOR CONTRIBUTIONS

Conceptualization: D.P., E.M., W.R.S., and F.C.

Data collection and curation: D.P., E.M., E.N., Q.L., T.A.R., L.E.D.V., K.T., and N.S.

Investigation and validation: D.P., E.M., E.N., Q.L., T.A.R., and L.E.D.V.

Writing, reviewing, and editing: D.P., E.M., W.R.S., and F.C.

Supervision and funding acquisition: F.C.

