## Supplemental Tables and Figures for "OocystMeter, a machine-learning algorithm to count and measure *Plasmodium* oocysts, reveals clustering patterns in the *Anopheles* midgut"

**SUPPLEMENTARY FIGURES AND TABLES**

### Table S1

a

| Oocyst-Meter iterations | Performance | | Number of infected midguts used | | Hyper-parameter tuning |
|---|---|---|---|---|---|
| | Average precision | Average recall | Training | Validation | |
| v1 | 83.0% | 86.2% | 69 | 17 | loss type, batch size, augmentation |
| v2 | 88.7% | 85.9% | 185 | 46 | loss type, batch size, augmentation |
| v3 | 94.4% | 90.0% | 185 | 46 | loss type, batch size, augmentation, proposal region number, positive fraction for evaluation |

b

| Midgut model | Performance | | Number of infected midguts used | | Hyper-parameter tuning |
|---|---|---|---|---|---|
| | Average precision | Average recall | Training | Validation | |
| v1 | 100.0% | 100.0% | 185 | 46 | loss type, batch size, augmenation, proposal region number, positive fraction for evaluation |

**Table S1. Average precision (AP) and average recall (AR) of the OocystMeter model using different numbers of training images and hyper-parameter tuning schemes.** Precision = correctly predicted positive / all predicted positives; Recall (sensitivity) = correctly predicted positive / all true positives. Intersect-over-union (IoU) = 0.7 was used as the threshold to calculate the AP and AR listed here. The OocystMeter model (v3) with AP = 94.4% and AR = 90.0% was used in subsequent analysis.

Figure S1

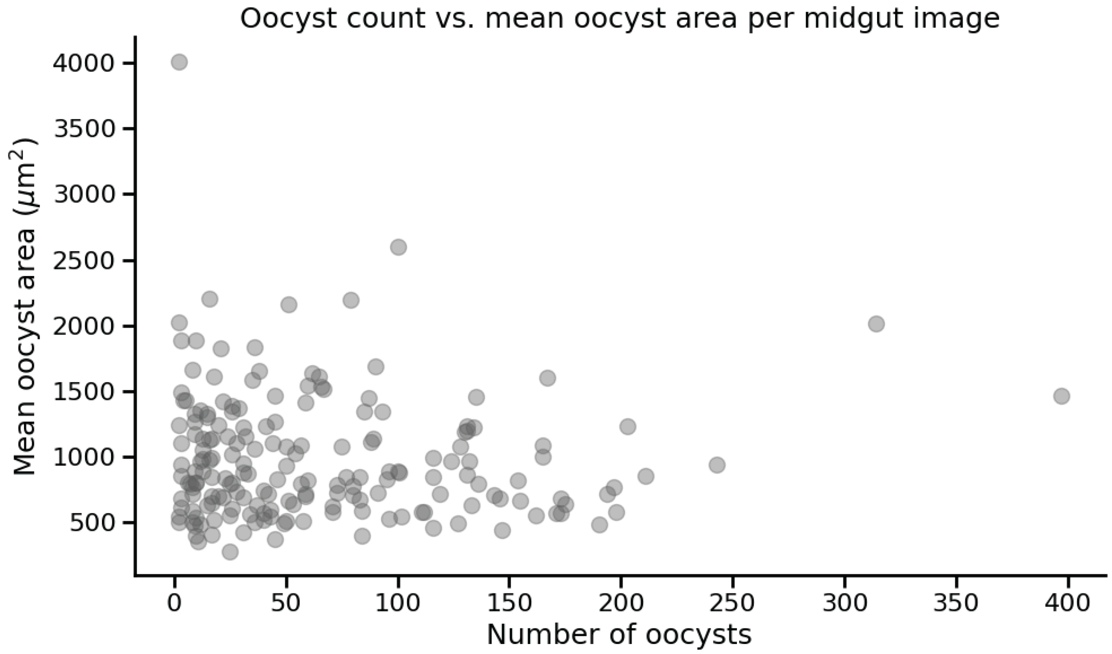


**Figure S1.** Scatter plot of oocyst count versus mean oocyst area for each midgut image in the training dataset.

### Figure S2

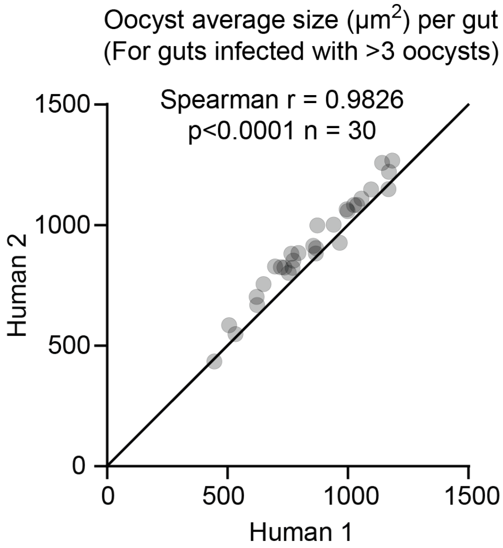


**Figure S2.** Systematic variation of oocyst size measurements between two humans. Each data point represents the average oocyst size in a midgut. The diagonal line is drawn in solid black.

### Figure S3

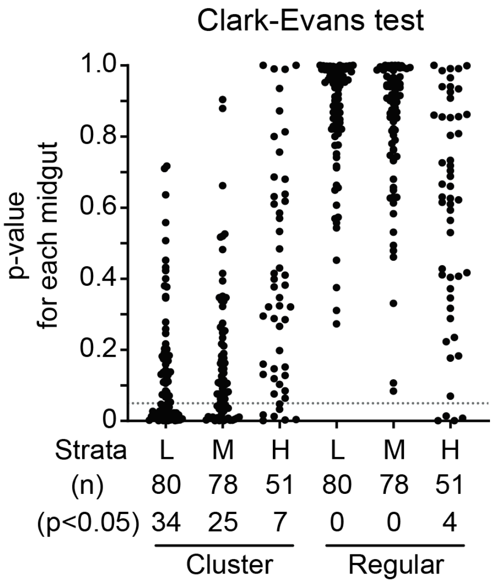


**Figure S3.** The Clark-Evans test was performed with "Cluster" or "Regular" alternative hypotheses (HA), $H_0$= Homogeneous Poisson point process (CSR). Each midgut generates a single multi-test-corrected p-value for each HA (significance level = 0.05).

### Figure S4

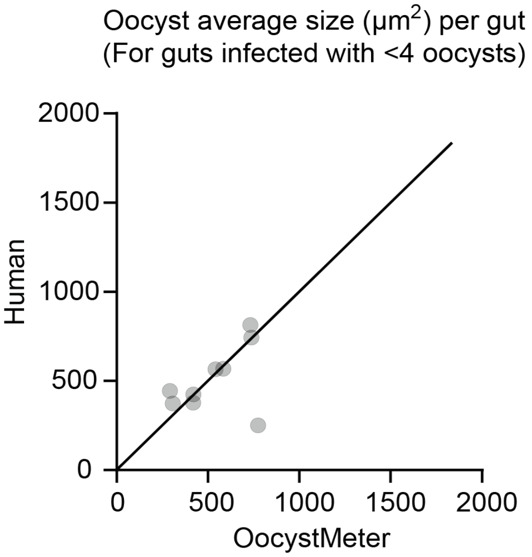


**Figure S4.** Average oocyst size measured by a human and the OocystMeter model for midguts infected with less than four oocysts. Each data point represents the average oocyst size in a midgut. The diagonal line is drawn in solid black.
